# SCAR and the Arp2/3 complex polarise the actomyosin cortex and plasma membrane organization in asymmetrically dividing neuroblasts

**DOI:** 10.1101/2023.01.05.522888

**Authors:** Giulia Cazzagon, Chantal Roubinet, Buzz Baum

## Abstract

While the Formin-nucleated actomyosin cortex has been shown to drive the changes in cell shape that accompany cell division in both symmetric and asymmetric cell divisions, it is not clear whether or not Arp2/3-nucleated branched actin filament networks also play a role. In order to look for mitotic roles of the Arp2/3 complex, here we use *Drosophila* neural stem cells as a model system. These cells are unusual in that they divide asymmetrically to produce a large and small daughter cell with different fates. Our analysis identifies a pool of Arp2/3-dependent actin-based membrane protrusions that form at the apical cortex of these cells as they enter mitosis. Strikingly, at metaphase, these protrusions co-localise with components of the SCAR complex. By perturbing Arp2/3 complex activity we show that this apical pool of actin likely functions to limit the accumulation of apical Myosin in metaphase. Following the onset of anaphase, the loss of these SCAR and Arp2/3 dependent structures then leads to a delay in the clearance of apical Myosin and to cortical instability at cytokinesis. These data point to a role for a polarised branched actin filament network in fine tuning the apical actomyosin cortex to enable the precise control of cell shape during asymmetric cell division.

## Introduction

While the Formin-nucleated actomyosin cortex is known to control the changes in cell shape that accompany division, much remains to be discovered about the role of branched actin networks during this process. The Arp2/3 complex is a nucleator of actin branched filaments, best known for its role in the formation of lamellipodial protrusions during adherent cell spreading and migration, in intracellular motility of pathogens, and in the fission of membranes during trafficking (Derivery et al., 2009; Kunda et al., 2003; Pollard, 2007; Rotty et al., 2013; Stevens et al., 2006).

It has been previously suggested that the Arp2/3 complex is mostly active in interphase, playing limited roles during mitotic entry and mitotic exit (Ramkumar & Baum, 2016). As examples of this, Arp2/3 dependent actin filaments have been shown to form at the centrosomes in cells entering and exiting mitosis and at the interface between newly divided cells (Farina et al., 2019; Herszterg et al., 2013; Plessner et al., 2019; Rajan et al., 2009; Trylinski & Schweisguth, 2019). This may be important since, when hyperactivated in patient cells, Arp2/3 dependent actin filament formation can impair chromosome segregation (Moulding et al., 2012).

However, a growing body of work carried out using mammalian cells in culture has also suggested that the Arp2/3 complex can generate actin filaments during mitosis. Indeed, in HeLa cells the Arp2/3 complex was shown to induce the formation of a rotating wave of actin filaments (Fink et al., 2011; Mitsushima et al., 2010) – although the role of this actin remains far from clear. In addition, the Arp2/3 complex has been implicated in the stabilisation of the mitotic cell cortex (Bovellan et al., 2014; Cao et al., 2020). However, it remains unclear whether or not the Arp2/3 complex plays a general function in mitosis. In addition, it remains to be tested whether or not Arp2/3 dependent actin filament formation plays important roles in the context of asymmetric cell division.

In general, the mechanisms that lead to shape changes in dividing *Drosophila* cells are very similar to those operating in vertebrate cells. In brief, upon entry into mitosis, the activation of Ect2 triggers Formin-dependent actin filament formation along with non-muscle Myosin II activation (hereafter called Myosin) to generate a contractile mitotic actomyosin cortex, which drives mitotic rounding (D’Avino et al., 2015; Matthews et al., 2012; Ramkumar & Baum, 2016; Rosa et al., 2015). Then at mitotic exit, cues from the spindle midzone and, more controversially, the anaphase chromatin polarise the mitotic cell cortex to allow the formation of a contractile actomyosin ring and division (D’Avino et al., 2015; Kiyomitsu & Cheeseman, 2012; Ramkumar & Baum, 2016; Rodrigues et al., 2015). However, there is currently little evidence for a role for the Arp2/3 complex in this type of cortical remodelling (Trylinski & Schweisguth, 2019).

Cells undergoing asymmetric divisions, like *Drosophila* neuroblasts, are likely to face additional challenges as they divide. At each round of asymmetric division, these neuronal stem cells produce two daughter cells with different size and fate: a big cell which retains stem cell feature, and a small cell, called ganglion mother cell (GMC), which divides again and differentiate into neurons or glial cells (Bello et al., 2008; Boone & Doe, 2008). In this system, polar cortical cues along with a polarised spindle function to break the symmetry of the mitotic cortex. As a result, at the onset of anaphase, Myosin is cleared from the apical cell cortex before being cleared from the basal cortex. This leads to biased cortical expansion, and to polarised cortical Myosin flows that drive the asymmetric positioning of the division ring and asymmetric division generating unequal sized sibling cells (Cabernard et al., 2010; Connell et al., 2011; Roubinet et al., 2017).

These actomyosin flows are coupled to membrane flows (Hannaford et al., 2018; LaFoya & Prehoda, 2021; Oon & Prehoda, 2019, 2021). In neuroblasts, membrane flows were recently shown to become polarized in early stages of mitosis, when they move apically. They are then reversed at the onset of anaphase, leading to the dispersal of the membrane domains across the cell surface (LaFoya & Prehoda, 2021). Interestingly, these movements have been shown to depend on cortical polarity and on the actomyosin cortex, implying a link between the membrane and the underlying cortex that could be important for asymmetric division (Hannaford et al., 2018; LaFoya & Prehoda, 2021; Oon & Prehoda, 2019, 2021).

In this paper we study the roles of the Arp2/3 complex and its upstream nucleation promoting actors in the regulation of membrane and cortical flows, mitotic cell shape, and division in fly neuroblasts. Using a combination of genetics and live cell imaging, our analysis reveals the existence of a pool of polarized actin-based membrane protrusions at the apical side of mitotic neuroblasts, which co-localise with SCAR complex components, whose organization depends on the Arp2/3 complex. We also show that this local remodelling of the actin cortex limits apical Myosin accumulation in metaphase, and when perturbed leads to cortical defects and membrane instability at cytokinesis. In this way, a local Arp2/3-dependent branched actin network appears to polarise the actomyosin cortex in mitotic neuroblasts to help guide the precisely choreographed cortical remodelling necessary for asymmetric cell division.

## Results

To study changes in membrane organization during passage through mitosis we began by using the Plekstrin Homology (PH) domain from the phospholipase CΔ1 (PLCΔ1) that interacts with the headgroup of the phosphatidylinositol 4,5-bisphosphate (PIP_2_) as a probe. This marker was chosen because it labels the plasma membrane and membrane protrusions, but not internal membranes (Figure 1A), and revealed changes in apical membrane organisation with mitotic progression (Figure 1A). To understand if these changes were local or global in nature, we compared PH intensity at the apical domain and the lateral membrane of mitotic cells. This analysis revealed that PH signal at the apical domain increases from prophase to metaphase, and then rapidly decreases upon the onset of anaphase, while the intensity of the signal in the lateral membrane remains constant (Figure 1B, and Supplemental Figure S1A-B). In these cells, an analysis of bright PH-labelled membrane domains also revealed a change in the polarity of membrane flows that depends on cell cycle stages, as previously reported (Supplemental Figure S1C) (LaFoya & Prehoda, 2021; Oon & Prehoda, 2019).

**Figure 1.**
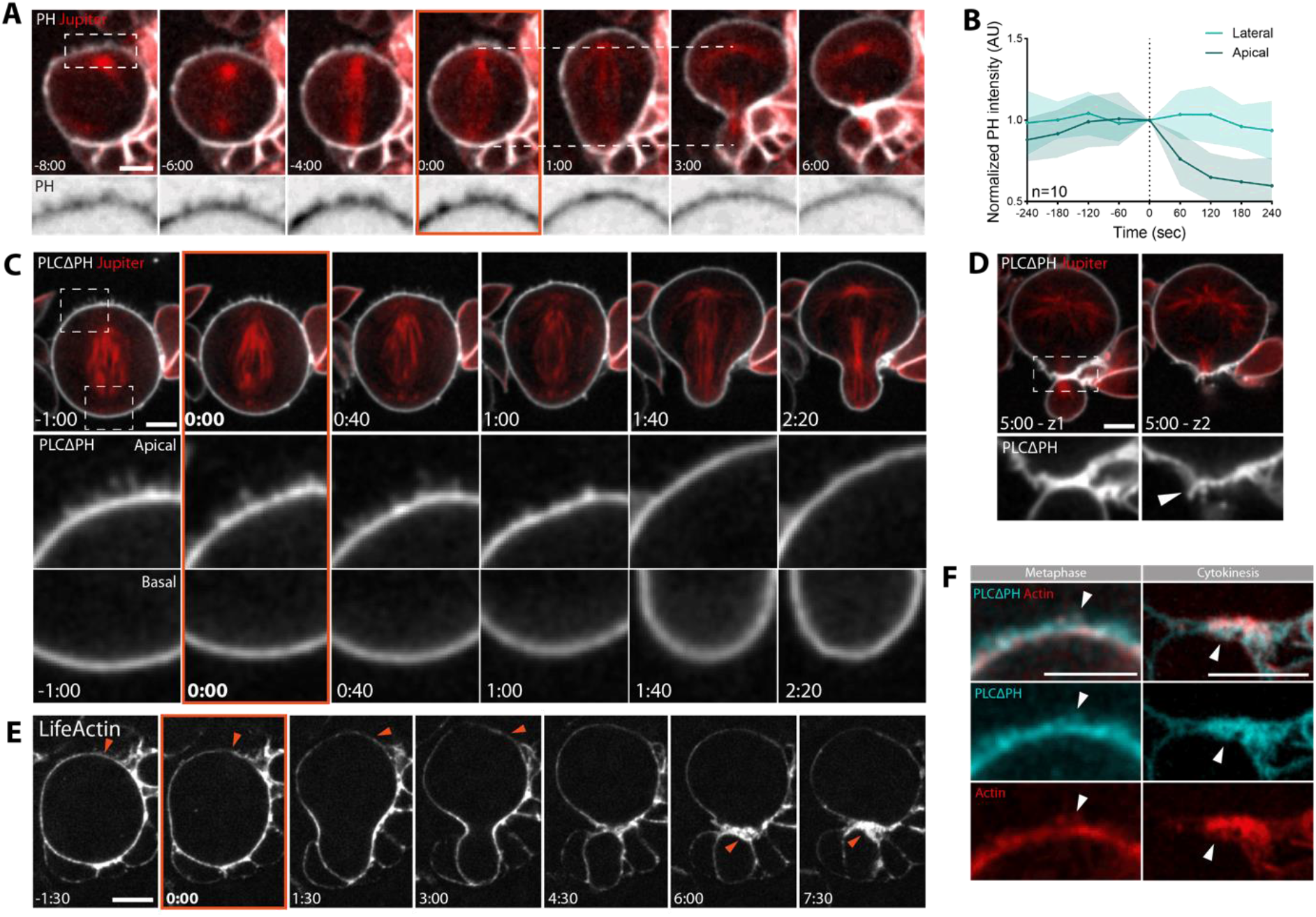
Dividing neuroblasts exhibit polarized membrane protrusions at metaphase, that disappear with cortical expansion following anaphase onset. **A**. Representative dividing neuroblast expressing membrane marker PLCΔPH::GFP and microtubule marker cherry::Jupiter, both expressed via wor-GAL4/UAS. Dotted lines highlight cortical expansion of the apical neuroblast domain, compared to the basal side that does not expand. **B**. Plot that shows PH intensity changes during mitosis progression in apical and lateral membrane. Data are included in Figure 1 – source data 1. **C**. Super-resolution imaging of neuroblast expressing PH::GFP and microtubule marker cherry::Jupiter. Inserts are apical and basal, respectively. **D**. Z-sections of cell at cytokinesis. Arrowhead points to membrane protrusion at the furrow. **E**. Dividing neuroblast expressing actin reporter UAS-LifeAct::GFP. Arrowheads point to actin being cleared from the apical cortex during metaphase-anaphase transition (−1:30 to 3:00 min), and to accumulation of actin at the cytokinetic furrow (6:00-7:00 min). **F**. Super-resolution live imaging of cells in metaphase or cytokinesis expressing PH::mCherry and LifeAct::GFP, both driven by wor-GAL4/UAS system. Arrowheads point to membrane protrusions that appear to be positive for actin filaments. Scale bar: 5 μm. Central and error bars: mean and SD.

To better understand these membrane dynamics, we used super-resolution spinning-disk confocal microscopy to image neuroblast metaphase-anaphase transition at higher temporal (20 sec/frame) and spatial resolution. In these movies, when the apical surface of cells was not in contact with overlying tissue, filopodia-like membrane structures could be seen forming at the apical cell surface at metaphase (Figure 1C, Apical insert, -1:00 to 0:40 min), while the basal cortex appeared relatively unchanged over this period (Figure 1C, Basal inserts). These apical protrusions were 0.7-1 μm in length, started to disappear 1 minute after anaphase onset, and were completely gone by the end of telophase (Figure 1C, Apical insert, 2:20 min); implying that they are absorbed as the apical cortex expands. The presence of polarized protrusions was confirmed using another membrane marker, GAP43 (Supplemental Figure S1D), demonstrating that they are a characteristic feature of the apical membrane independently of the reporter used. Sometime later, the membrane marker also revealed a population of protrusions forming between the two daughter cells at the site of cleavage (Firgure 1A and 1D, arrowhead), similar to those described previously at new cell interfaces in other cell types (Herszterg et al., 2013; Rajan et al., 2009; Trylinski et al., 2017).

Since membrane protrusions are often actin-based, we used the LifeAct::GFP probe to image the localisation of actin filaments in mitotic neuroblasts. This revealed an accumulation of actin at the apical cortex, which was cleared during the metaphase-anaphase transition (Figure 1E, arrowheads from -1:30 to 3:00 min). Once again, actin-rich protrusions were also observed later at the furrow following cytokinesis (Figure 1E, arrowheads 6:00-7:30 min). Both sets of actin structures were found to colocalise with the PH signal in flies carrying both markers (Figure 1F, arrowheads).

Since actin and membrane protrusions at the cytokinetic furrow in *Drosophila* sensory organ precursors (SOPs) have previously been shown to depend on the actin nuclear-promoting factor SCAR (Georgiou & Baum, 2010; Trylinski & Schweisguth, 2019), we decided to test if this was also the case in the neuroblast. To do so, we dissociated larval brains to look at isolated neuroblasts, and imaged cells expressing SCAR::GFP at high spatial resolution. This experiment revealed the asymmetric localization of SCAR at the apical side of the neuroblast in metaphase (Figure 2A, arrowheads, and Figure 2B). When the neuroblast entered anaphase, SCAR::GFP was lost from the apical cortex, before accumulating at the basal side of the cell sometime later, where it became concentrated at the cytokinetic furrow (Figure 2A, arrowhead, and Figure 2B). Since SCAR is part of a stable multiprotein complex, we validated this localization using a second complex component, Abi (Supplemental Figure S2). We then imaged SCAR and membrane protrusions in parallel in mitotic cells expressing both PH::mCherry and SCAR::GFP (Figure 2C). While the fluorescent signal was low and there were few protrusions at prophase (Figure 2C, -8:00 minutes). By metaphase (Figure 2C, -2:00 min), numerous SCAR-positive membrane rich protrusions were visible decorating the apical surface of neuroblasts (arrowheads). When this was quantified by measuring intensity on a line drawn along the apical cell surface, it was clear that the peaks of SCAR and PH exhibit partial overlap. At anaphase (Figure 2C, 1:00 minute), the PH signal appeared to smooth out, while SCAR signal was still visible in puncta on the cortex in places that lack protrusions. These data suggest a relatively tight correlation between the presence of SCAR signal and membrane protrusions in metaphase cells.

**Figure 2.**
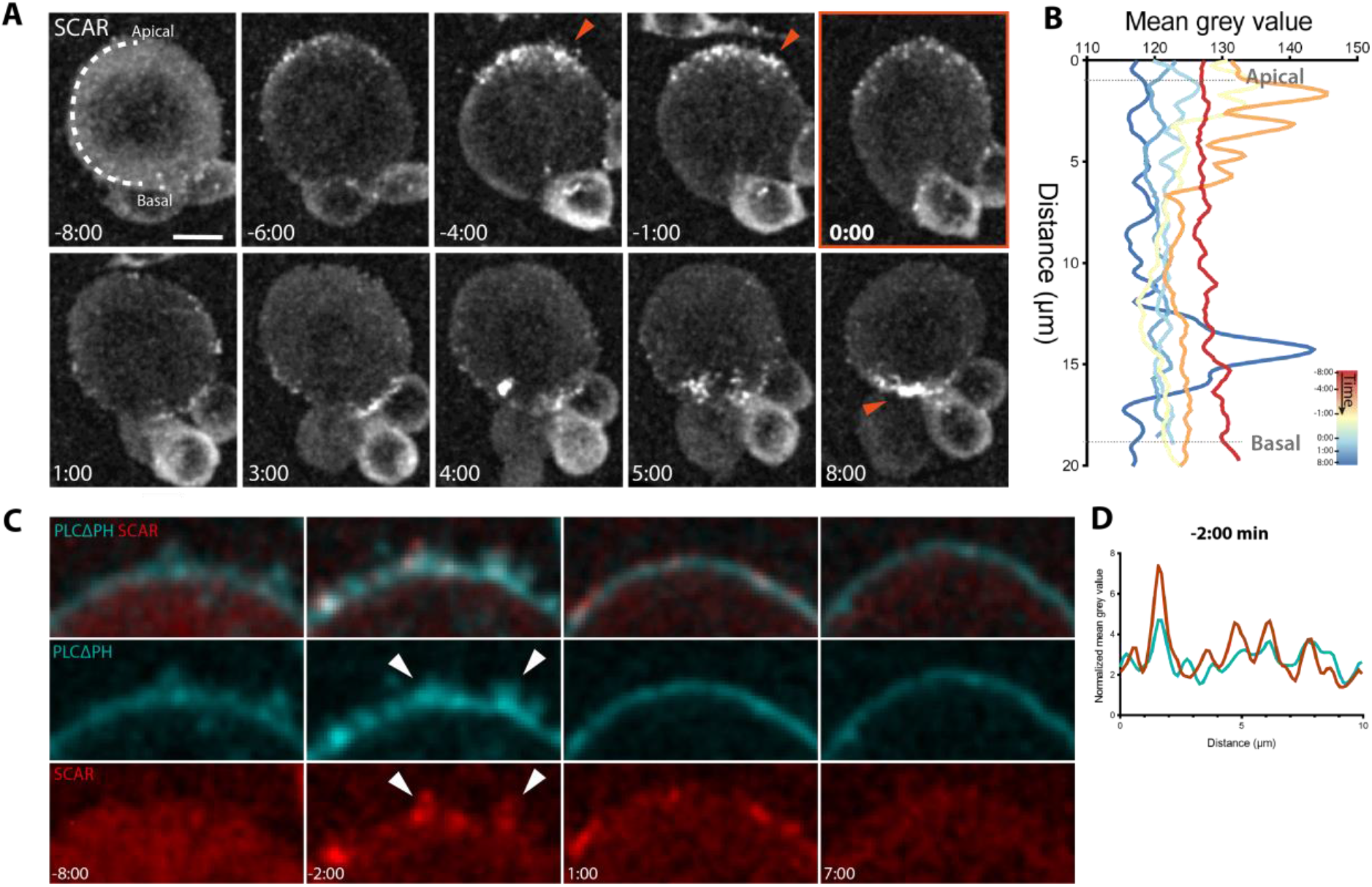
SCAR co-localizes with membrane protrusions in dividing neuroblasts. **A**. Representative super-resolution maximum z-projection of dissociated neuroblast expressing SCAR::GFP driven by wor-GAL4/UAS system. Arrowheads point to SCAR signal at the apical side of the neuroblast in metaphase and at the furrow in cytokinesis. **B**. Graph showing mean SCAR signal intensity, that was acquired by drawing a line from apical to basal side of the cortex, like depicted at time -8:00 in A. Individual intensities are color-coded by time. **C**. Super-resolution imaging of a representative neuroblast, showing partial co-localization of membrane protrusions (UAS-PH::mCherry) and SCAR signal (UAS-SCAR::GFP). **D**. The graph was obtained by drawing a line in the portion of cortex included in the insert at -2:00 minutes from anaphase onset and averaging PH or SCAR signals. Data were normalized by subtracting background. Scale bar: 5 μm.

To test whether apical protrusion formation in this system depends on the SCAR and Arp2/3-dependent nucleation of branched actin filaments, as suggested by these data, we treated neuroblasts expressing the PH membrane marker with Arp2/3 inhibitor CK-666, and then imaged the treated cells at high resolution. While CK-666 treated cells possessed PH-rich membrane domains in metaphase like those seen in the control, the small molecule had a profound impact on their organisation (Figure 3A and 3B). In Arp2/3-inhibited cells, the excess membrane was observed forming small rounded structures (Figure 3B, arrowhead), rather than filopodia-like protrusions (Figure 3A). Thus, while the Arp2/3 complex activity is not required for the accumulation of membrane apically, it is required for its proper organisation. During the metaphase-anaphase transition, the membrane domains present in CK666-treated cells smoothed out so that they were no longer visible by cytokinesis (Figure 3B’) with similar kinetics to the loss of membrane protrusions in the control (Figure 3A’). Thus, membrane flows during metaphase-anaphase transition are not abolished by inhibition of the Arp2/3 complex (Supplemental Figure S3A).

**Figure 3.**
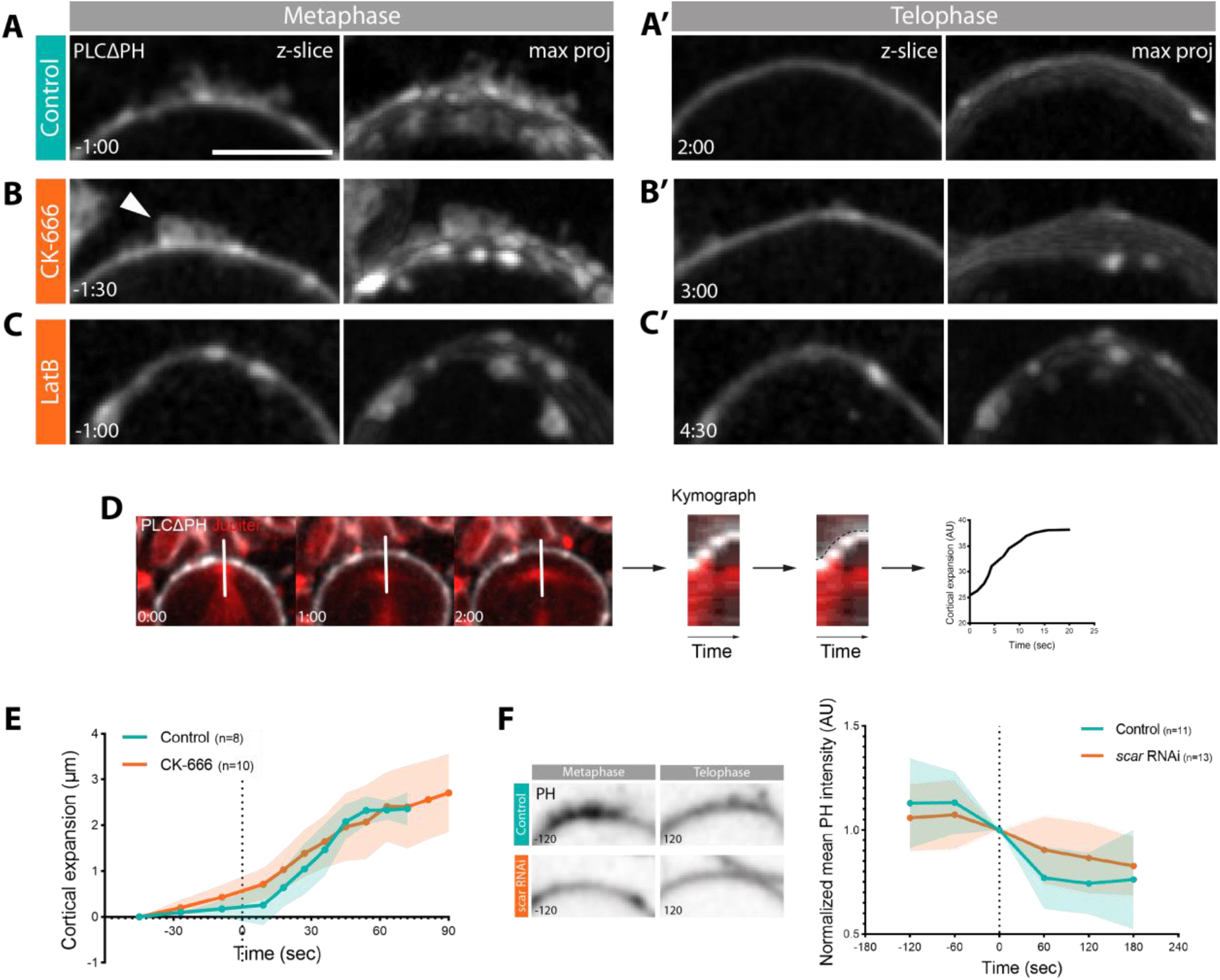
The Arp2/3 complex is required for the organization of apical actin-rich protrusions and for the precise dynamics of apical expansion. **A-C’**. Inserts depicting z-slice and maximum intensity z-projection of apical side of metaphase and telophase neuroblasts expressing PH::GFP marker. Image shows state of the cortex in control (**A-A’**) and the effect of CK-666 (**B-B’**) and Latrunculin B (**C-C’**). **D**. Schematics showing how plot in E was obtained: kymographs were generated drawing a line like shown in figure, the movement of the membrane was manually traced and coordinates were exported and plotted. The final result is represented by the graph on the right. **E**. Graph showing mean of membrane expansion during anaphase, for control (CK-689) and CK-666 treated cells. Coordinates were centred to start at (x=0, y=0) and to plot the mean, a linear interpolation of the x set of coordinates was performed. The slope of the two curves is significantly different (t-test, P-value=0.0044). Data are included in Figure 3 – source data 1. **F**. Images of neuroblasts apical cortex and plot showing changes in PH intensity during time in control and scar-dsRNA cells. 2-way ANOVA comparison between control and RNAi: *P ≤ 0.05. Data are included in Figure 3 – source data 2. Scale bar: 5 μm. Central and error bars: mean and SD.

To determine more generally the role of actin filament in the formation of the polarized membrane protrusions in this system, we also treated cells with Latrunculin B (LatB), which binds to monomers leading to the rapid loss of actin filaments (Morton et al., 2000). Again, local patches of apical membrane were seen becoming enriched in metaphase in these drug-treated cells. However, these seemed disorganised and appeared to protrude into the cell (Figure 3C), instead of forming spike-like outward-facing protrusions like those seen in control cells (Figure 3A). These disorganised patches of membrane were still visible in later stages of mitosis (Figure 3C’), implying that LatB blocks all cortical remodelling.

To test if inhibiting the Arp2/3 complex also has an effect on cortical dynamics that take place during metaphase-anaphase transition, we extracted the coordinates of the movement of the membrane during the cortical expansion phase (Figure 3D). The data for individual cells was then plotted and averaged (Figure S3B and 3E). In control cells, the resulting curve depicting the dynamics of apical expansion had a clear sigmoid shape, due to a sudden movement that quickly came to a stop (Figure 3D, S3B-C). By contrast, in cells in which Arp2/3 was inhibited by CK-666, apical expansion was slower and linear (Figure 3D, S3B-C). When the slopes of the two curves were computed by fitting a sigmoid and compared, they were found to be significantly different (t-test, P-value=0.0044). Thus, although the treatment does not block apical expansion, it changes the dynamics of cell shape changes following the onset of anaphase.

To confirm the role of SCAR in regulating membrane flows at metaphase-anaphase transition, we measured again PH intensity over the same time interval in cells where SCAR was inhibited through RNAi. While the PH probe accumulated apically before anaphase onset in control cells, and then decreased during cortical expansion (Figure 3F) as expected (Figure 1B), the PH signal remained low and changed little as SCAR RNAi cells underwent cortical expansion. Indeed, there was a significant difference between the two curves (2-way ANOVA, P-value=0.0447) (Figure 3F). These data suggest that SCAR likely works together with the Arp2/3 complex in regulating organization of the apical cortex in mitotic neuroblasts.

The asymmetric cortical expansion observed in neuroblasts is thought to reflect a difference in the timing of apical and basal Myosin clearance at anaphase (Connell et al., 2011; Roubinet et al., 2017). Since SCAR is concentrated at the apical cell cortex, which is the first to lose Myosin at the onset of anaphase to trigger an apico-basal directed cortical flow (Supplemental Figure S4A), we decided to test if the Arp2/3 complex has a role in this process. To do so, we imaged non-muscle Myosin II, using the Sqh::GFP reporter, at higher temporal (15 sec/frame) and spatial resolution (Figure 4A). We then quantified Myosin levels at both the apical and basal sides of *arp3* mutant neuroblasts, and used the heterozygous as control. In heterozygous animals, cortical flow resembles that seen in the wild type (Figure 4B, and Supplemental Figure S4B), with the clearance of apical Myosin beginning around 15 seconds after anaphase onset, followed by basal clearance at around 60 seconds (Figure 4B, arrowheads). By contrast, in homozygous *arp3* mutant animals there was a significant 15 second difference in the timing of Myosin clearance from both the apical and basal cortex following the onset of anaphase. This occurred at 30 seconds for the apical cortex, and 90 seconds at the basal side (Figure 4B’, arrowheads, and 4B”).

**Figure 4.**
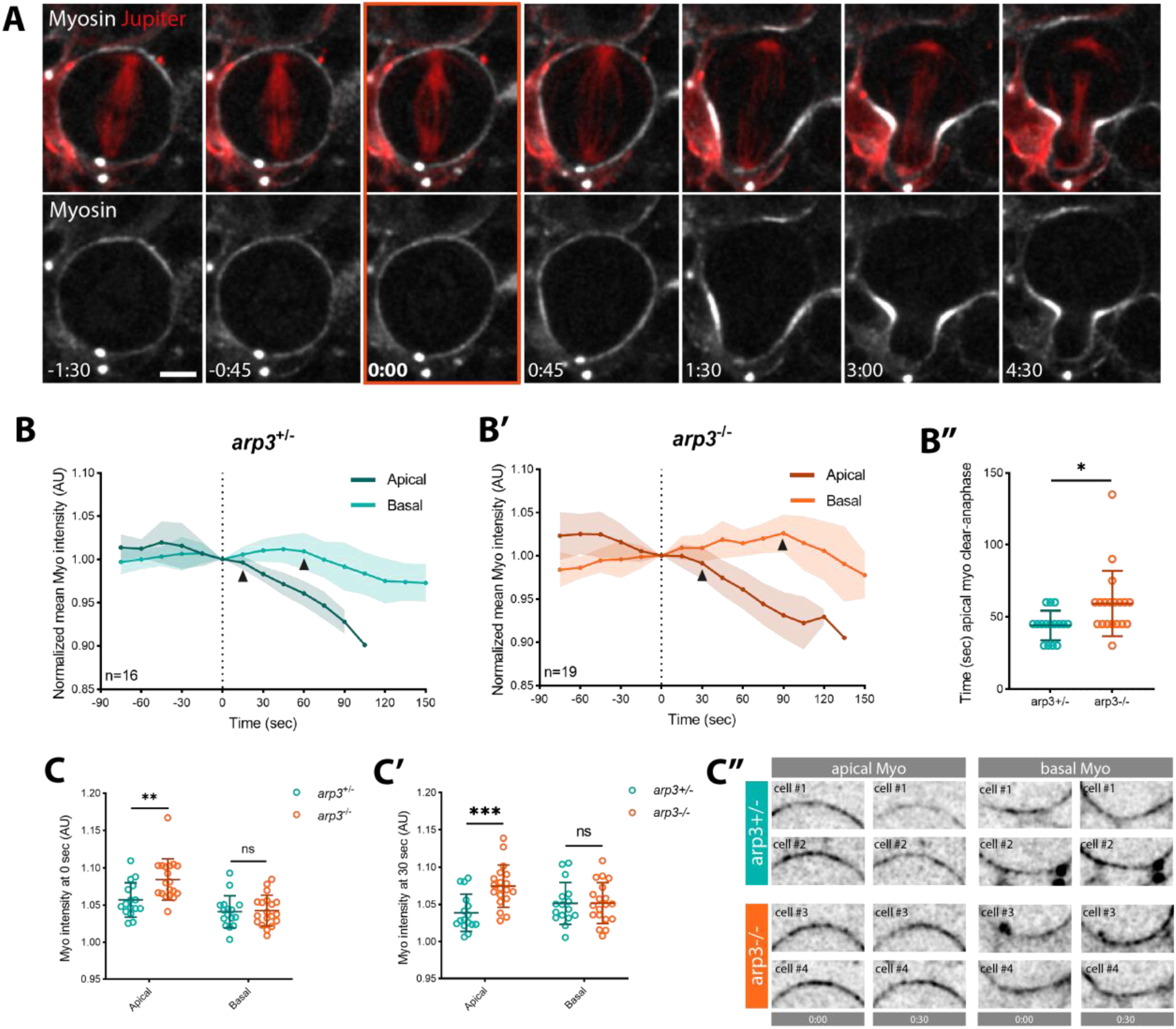
Arp2/3 modulates apical Myosin dynamics in mitotic neuroblasts. **A**. Super-resolution time-lapse image of dividing neuroblast expressing non-muscle Myosin II marker (Sqh::GFP) and microtubule marker (UAS-cherry::Jupiter). **B-B’**. Graphs showing Myosin intensity changes during metaphase-anaphase transition in heterozygous *arp3*^*+/-*^ (arp3^EP3640^/TM6B) (**B**) and mutant *arp3*^*-/*-^ (arp3^EP3640^/Deficiency) (**B’**) expressing Sqh::GFP and UAS-cherry::Jupiter. Arrowheads mark the start of Myosin clearance. Myosin intensity was normalized by subtracting the background and centred at time 0. **B”**. Plot showing time of apical Myosin clearance relative to anaphase onset (t-test). **C-C’**. Myosin intensity at time 0 (**C**) and 30 seconds (**C’**) at the apical and basal sides, compared between *arp3*^*+/-*^ and *arp3*^*-/-*^ (2-way ANOVA). Myosin intensity was normalized by subtracting background. **C”**. Examples of apical and basal cortex with Myosin signal at time 0 and 30 seconds after anaphase onset for arp3+/- and arp3-/-. Asterisks denote statistical significance. ns, not significant P > 0.05, *P ≤ 0.05, **P ≤ 0.01, ***P ≤ 0.001. Data are included in Figure 3 – source data 2. Scale bar: 5 μm. Central and error bars: mean and SD.

Furthermore, when comparing Myosin intensities at the onset of anaphase and 30 seconds later, it became clear that *arp3* mutant cells accumulate higher levels of apical cortical Myosin than their control counterparts (Figure 4C-C”). This strongly suggests that the assembly of a branched SCAR and Arp2/3 dependent actin network negatively regulates the assembly of cortical Myosin, to promote rapid apical Myosin clearance at the onset of anaphase.

To determine the effect of SCAR/Arp2/3 activity on the neuroblast division more generally and to understand how these subtle but significant changes in cortical remodeling impact later stages of mitosis, we imaged a large number of neuroblasts expressing Myosin marker Sqh::GFP and the microtubule marker UAS-cherry::Jupiter (Figure 5A). To perturb Arp2/3 activity, we then used Arp2/3 inhibitor CK-666 and genetic tools (mutation and RNAi-mediated silencing of *arp3* (Figure 5A’-B’, 5D, and Supplemental Figure S5A). While similar phenotypes were observed in all cases, the strongest was observed following the chemical inhibition of Arp2/3. In CK-666 treated cells, but not control cells, Myosin was observed ectopically accumulating at the cortex after the completion of cytokinesis, leading to an aberrant late constriction of the plasma membrane, which generated a large rounded protrusion (Figure 5A’, arrowheads and 5D). However, the necks of these protrusions were never seen closing. A milder version this phenotype was observed in cells homozygous for *arp3* mutations and in *arp3* RNAi cells, where the accumulation of ectopic Myosin was accompanied by a range of cortical defects (Figure 5B-B’, 5D, and Supplemental Figure S5A). Finally, we tested whether or not similar defects were in somatic *scar* mutant clones in the larval brain and in cells in which *scar* was knocked-down with RNAi (Figure 5C-C’, 5D and Supplemental Figure S5B). These cells depleted for SCAR activity exhibited ectopic Myosin localization and cortical defects after the completion of cytokinesis similar to those observed following perturbation of the Arp2/3 complex (Figure 5C’ arrowheads and Supplemental Figure S5B arrowheads). The cell contour overlays on the side of the montages show how the membrane changes over time in cells with conditions compared to controls (Figure 5A-C’). These experiments confirm a role for the SCAR and Arp2/3 complexes in regulating proper asymmetric neuroblast division, and show that dysregulation of one of these proteins leads to a various range of defects from metaphase until the end of cytokinesis.

**Figure 5.**
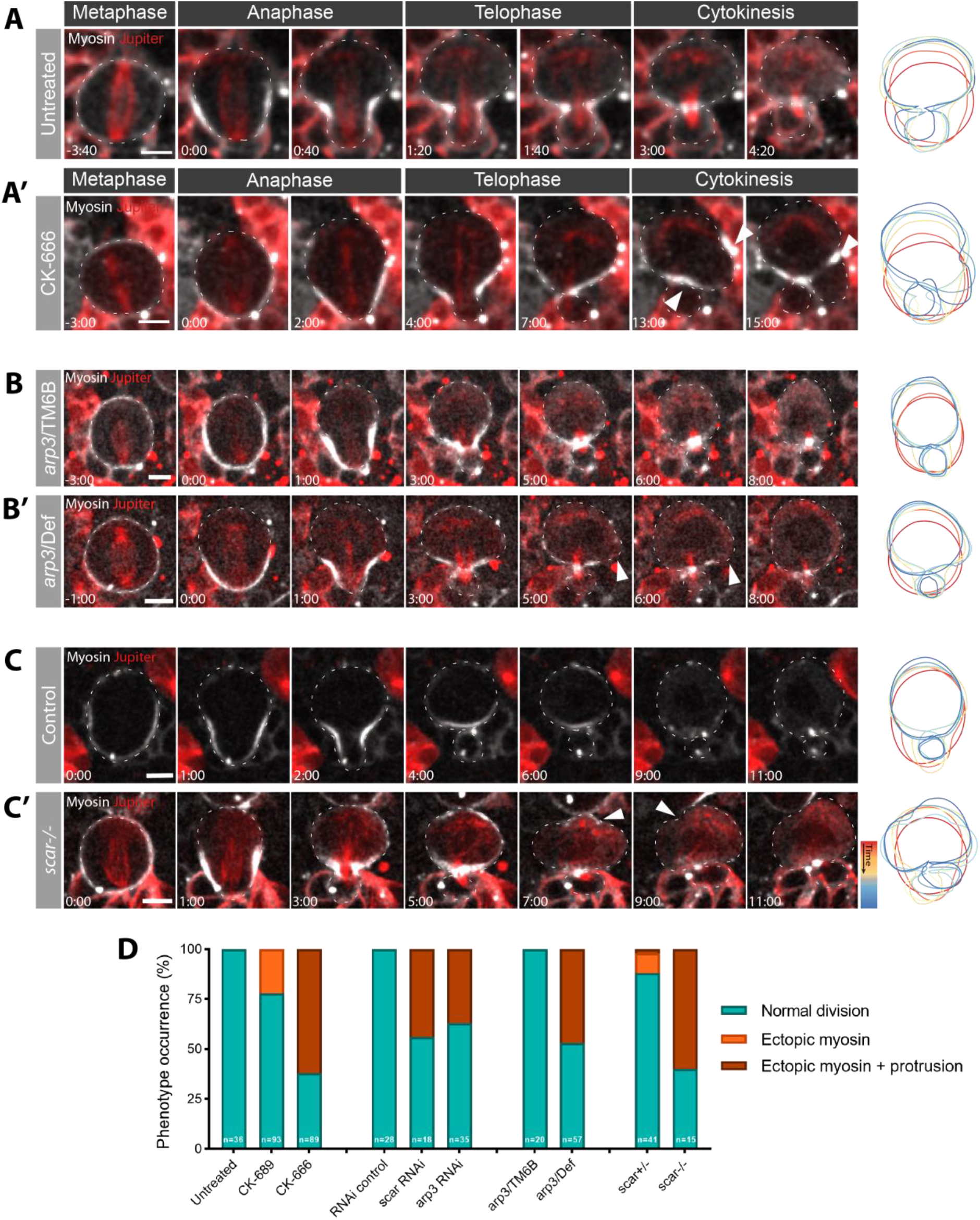
SCAR or Arp2/3 inhibition in the dividing neuroblast leads to cortical instability after cytokinesis. **A-C’**. Time-lapse images of dividing neuroblasts expressing a non-muscle Myosin II reporter Sqh:GFP and a microtubule marker UAS-cherry::Jupiter. Examples of untreated (**A**) and CK-666 treated (**A’**) neuroblasts. Mutant *arp3* (arp3/Deficiency) (**B’**) and control (**B**, heterozygous arp3/TM6B). Neuroblast in *scar* mutant clone (**C’**, *scar-/-*) and control cell (**C**). Arrowheads point to cortical defects and membrane protrusions due to Arp2/3 or SCAR inhibition. Cell contours on the right show for each condition how cell shape changes from anaphase to cytokinesis. Contours are color-coded by time. **D**. Graph showing percentages of cells affected by specific phenotype in each set of conditions. Scale bar: 5 μm.

## Discussion

In this study we present a novel role for SCAR and the Arp2/3 complex in regulating membrane organization and cell shape changes in metaphase-anaphase transition in neural stem cells in the fly. Thanks to high-resolution spinning-disk microscopy we were able to image the asymmetric accumulation of filopodia-like membrane protrusions rich in SCAR, actin and PIP_2_ lipids at the apical cortex of these cells in metaphase (Figure 1). Although Arp2/3 and SCAR are usually associated with the formation of lamellipodia, this is not unexpected, since in flies these proteins have been shown to generate filopodial-like cell extensions from an underlying branched actin network in cells in culture and *in vivo* (Biyasheva et al., 2004; Georgiou & Baum, 2010).

While the Arp2/3 complex is responsible for protrusion formation, we still observed an accumulation of excess apical membrane in patches following perturbations in Arp2/3 complex activity (Figure 3). This suggests that although the Arp2/3 complex nucleates a branched-actin network which might create the scaffold for membrane protrusion formation, it is not necessary for apical membrane accumulation (although the Latrunculin experiment suggests that actin filaments may play a role in this process).

The apical, actin-rich membrane protrusions formed as cells enter mitosis were quickly absorbed as cells underwent cortical expansion as they entered anaphase (Figure 1). While this might lead to the suggestion that protrusions provide a pool of excess membrane to facilitate cortical expansion, inhibiting the Arp2/3 complex and protrusion formation did not block the accumulation of an apical pool of membrane rich in PIP_2_ and did not prevent the cell from completing division (Figure 5; Supplemental Figure S3). Thus, the protrusions themselves do not appear to act as a functionally important membrane reservoir.

On the other hand, the lack of Arp2/3 activity does alter the dynamics of cortical expansion, cortical stability, and leads to aberrant changes in the shape of cells undergoing cytokinesis (Figure 3 and Figure 5). These data point to a role for the complex in polarising the cortex to regulate relaxation of the apical pole and the dynamics of shape changes that follow. In this process, the SCAR complex would provide a cue to bias the accumulation of Arp2/3 leading to the formation of a branched apical actin network.

The inhibition of Arp2/3 complex activity also leads to an increase of apical Myosin at the onset of anaphase, without affecting the basal Myosin pool, and leads to a delay in the clearance of apical Myosin relative to the onset of anaphase (Figure 4). Interestingly, in *Drosophila* salivary glands, where the actomyosin cortex is used to collapse large spherical secretory vesicles, the Arp2/3 complex has been proposed to form stripes of branched actin that help to break the symmetry of the Formin-nucleated actomyosin cortex around the vesicle to facilitate the collapse (Rousso et al., 2016). We propose that SCAR and Arp2/3 complexes act in a similar way to pattern the actomyosin cortex of mitotic neuroblasts to facilitate asymmetric division. In this case, an apical Arp2/3 dependent actin network may limit the apical accumulation of Myosin (Muresan et al., 2022; Truong Quang et al., 2021) to facilitate rapid apical cortical expansion at the onset of anaphase. While it is not clear precisely how these early changes in cortical remodelling dynamics affect cytokinesis, it is possible that defects early on in the process lead to stronger phenotypes at later stages of division. Nevertheless, it remains possible that the loss of branched actin from the apical cortex has other effects on the system that alter cortical instability at later stages in other ways.

Our data suggest that SCAR functions as the main Arp2/3 activator in the generation of apical membrane protrusions in this system, since reduction in SCAR (using mutants and RNAi) leads to the same class of phenotypes as those observed following reduction in Arp2/3 activity (drugs, mutants and RNAi) (Figure 5). While this is the case, the membrane constriction phenotype observed at cytokinesis is stronger and more consistent in CK-666 treated cells than it is following perturbations of SCAR function. Since flies possess additional NPFs, it is also possible that WASH and WASp play minor roles as Arp2/3 activators in this system. This is because, while there is often a clear separation between Arp2/3 nucleation promoting factors, with SCAR being responsible for lamellipodia formation, WASp being involved in filopodia, and WASH being involved in trafficking, this is not always the case (Campellone & Welch, 2010; Chesarone & Goode, 2009). If they have partially redundant roles, this may explain why the Arp2/3 phenotypes tend to be stronger than those seen following reductions in SCAR levels.

At the same time, the SCAR depletion appears clear and fits that described by previous studies that implicated SCAR in the formation of thin actin-based protrusions (Georgiou & Baum, 2010; Trylinski & Schweisguth, 2019; Zallen et al., 2002). Moreover, SCAR appears to be in the right place at the right time to generate apically polarised Arp2/3 dependent protrusions. While it is not clear from our work how SCAR is localised, its polarized localization resembles that of aPKC and Par3 (Loyer & Januschke, 2020; Petronczki & Knoblich, 2000; Wodarz et al., 2000). In both *Drosophila* neuroblasts and *C. elegans* a link between these proteins and F-actin, Myosin and membrane domains has been clearly established (LaFoya & Prehoda, 2021; Oon & Prehoda, 2021; Scholze et al., 2018). Future work will be necessary to elucidate whether SCAR is recruited by the polarity pathway and the mechanisms involved. In general, however, our data shows how the polarised localisation of SCAR locally activates Arp2/3 to break the symmetry of the cortical actomyosin network in metaphase. At the onset of anaphase, the presence of this apical branched actin network may then help to tune the actomyosin cortex to enable the precise control of changes in cell shape and membrane organisation required for asymmetric cell division.

## Methods

### Fly strains and genetics

Mutant chromosomes were balanced over Cyo::ActGFP or TM6B, Tb. The following mutant alleles and RNAi lines were used: Arp3^EP3640^ (BL17149, Bloogminton) (Rørth, 1996), Df(3L)Exel6112 (removes Arp3, BL7591, Bloogminton), SCAR^Δ37^ FRT40A (BL8754, Bloomington) (Zallen et al., 2002), SCAR RNAi (BL36121, Bloomington).

### Transgenes and fluorescent markers

Sqh::GFP (Royou et al., 2002), and UAS-cherry::Jupiter (Cabernard & Doe, 2009) from C. Roubinet. UAS-PLCΔPH::GFP (BL39693, Bloomington), UAS-PH::mCherry (BL51658, Bloomington), UAS-LifeAct::GFP (BL58718, Bloomington), UAS-SCAR::GFP (from M. González-Gaitán). Transgenes were expressed using the neuroblast-specific driver worniu-Gal4 (Albertson & Doe, 2003).

### Live imaging sample preparation

Larvae were dissected to extract the brains in imaging medium (Schneider’s insect medium mixed with 10% FBS (Sigma), 2% PenStrepNeo (Sigma), 0.02 mg/mL insulin (Sigma), 20mM L-glutamine (Sigma), 0.04 mg/mL L-glutathione reduced (Sigma) and 5 μg/mL 20-hydroxyecdysone (Sigma)). Brains were then transferred with the medium onto 15μ-slide angiogenesis (Ibidi), brain lobes facing down, and imaged. When brain dissociation was performed, larvae were dissected in Chan & Gehring solution 2% FBS (CG-FBS) to extract the brain (Chan & Gehring, 1971). GC-FBS composes as follow: NaCL 3.2 g/l, KCl 3 g/l, CaCl_2_-2H_2_O 0.69 g/l, MgSO_4_-7H_2_O 3.7 g/l, Tricine buffer Ph7 1.79 g/l, glucose 3.6 g/l, sucrose 17.1 g/l, BSA 1g/l and FBS 2%. Papain (Sigma, #P4762-50MG, 10 mg/ml) and collagenase (Sigma, #C2674-1G, 10 mg/ml) were added to the brains in CG-FBS solutions and they were incubated at 29°C for 45 minutes, to activate the enzymes. After incubation, brains were washed with imaging medium and finally dissociated through vigorous pipetting. The brains were then transferred with the medium onto 15μ-slide 8 well (Ibidi) and imaged.

### Imaging

Super-resolution imaging was performed on a CSU-W1 SoRa spinning disk confocal microscope (Nikon Ti Eclipse 2; Yokogawa CSU-W1 SoRa spinning disk scan head) with 60×/1.40 N.A oil objective and equipped with a photometrics prime 95B scientific CMOS camera. Whole brain live imaging was performed on a UltraView Vox spinning disk confocal microscope (Perkin Elmer Nikon TiE; Yokogawa CSU-X1 spinning disc scan head) with 60×/1.40 N.A oil objective and equipped with a Hamamatsu C9100-13 EMCCD camera. Whole brain imaging has been acquired with a z-stack spacing of 1 μm, while single cell imaging with a spacing of 0.7 μm. Time resolution was 60 seconds per frame, unless specified otherwise. Both microscopes are equipped with a temperature-controlled environment chamber set at 26° C for the experiments.

### Treatments

For chemical treatments to inhibit the Arp2/3 complex, the inhibitor CK-666 (Sigma #SML0006, final concentration 400 μM) or the inactive equivalent compound, CK-689 (Sigma #182517, final concentration 400 μM), were added before live imaging. To induce actin depolymerization, Latrunculin B (Sigma #L5288-1MG) at a final concentration of 10 μM was added to the media.

### Image processing

All image analysis was carried out on unprocessed raw images. For clarity, images displayed in this work were processed using ImageJ software (Schindelin et al., 2012). Background was (rolling ball radius 50 pixel) and a Gaussian Blur applied (radius 1). As stated in Figure legends, images represent a single confocal z-stack section or a maximum z-projection. In all figures, the time point 0 is anaphase onset, defined in this work as the first frame where the spindle starts to separate. Figures were assembled using Adobe Illustrator CS6.

### Image analysis

Experiments in which cortical PH or Myosin intensity were measured, a line of a specific width and length was drawn on the area of interest and the mean pixel value was calculated. The data were normalized by subtracting the background and were centred at the time 0. To calculate the movement of the membrane during cortical expansion, a maximum projection of 3 z-slices from the centre of cells dividing along the axis parallel to the field of view was generated. A line was drawn from the centre of the spindle to the apical membrane, starting two time-frames before anaphase onset. A kymograph was generated from this line, the movement of the membrane was traced and the set of coordinates were used to generate the curves in the graphs. Coordinates were centred to start at (x=0, y=0). To plot the mean, the curves were interpolated in Excel using linear interpolation. The mean curves ware fitted with a sigmoid curve in Prism. The slopes were calculated by fitting each curve with a sigmoid in Prism, the slopes of curves for control and treated cells were then compared with a t-test.

### Statistical analysis

For experiments with quantification, the data was collected from at least 2 independent experiments, and, for each independent experiment, at least 2 brain lobes were imaged. For the analysis, “n” refers to the number of cells analysed and is represented on the graph or mentioned in figure legends. Statistical significance was determined with Student’s t test where two groups were compared and 2-way ANOVA where more than two groups were compared, using GraphPad Prism 9 software. In all figures the Prism convention is used: ns (P > 0.05), *(P ≤ 0.05), **(P ≤ 0.01), ***(P ≤ 0.001) and ****(P ≤ 0.0001). In all graphs showing mean, the error bars correspond to standard deviation (SD).

## Funding

G.C. was supported by MRC (1621658). C.R. and B.B. were supported by a Cancer Research UK program grant (C1529/A17343). We thank the MRC LMB for their generous support.

## Author contributions

B.B. and G.C. conceived the study with input from C.R. Initial observations on cortical defects at cytokinesis after CK-666 treatment were made by C.R. G.C performed experiments and analysis. B.B and G.C. wrote the manuscript. C.R. provided advice.

## Acknowledgments

We thank Emmanuel Derivery and Guillaume Charras for reading the manuscript and providing comments.

## Competing interests

Authors declare that they have no competing interests.

## Supplementary figures

**Supplemental Figure S1.**
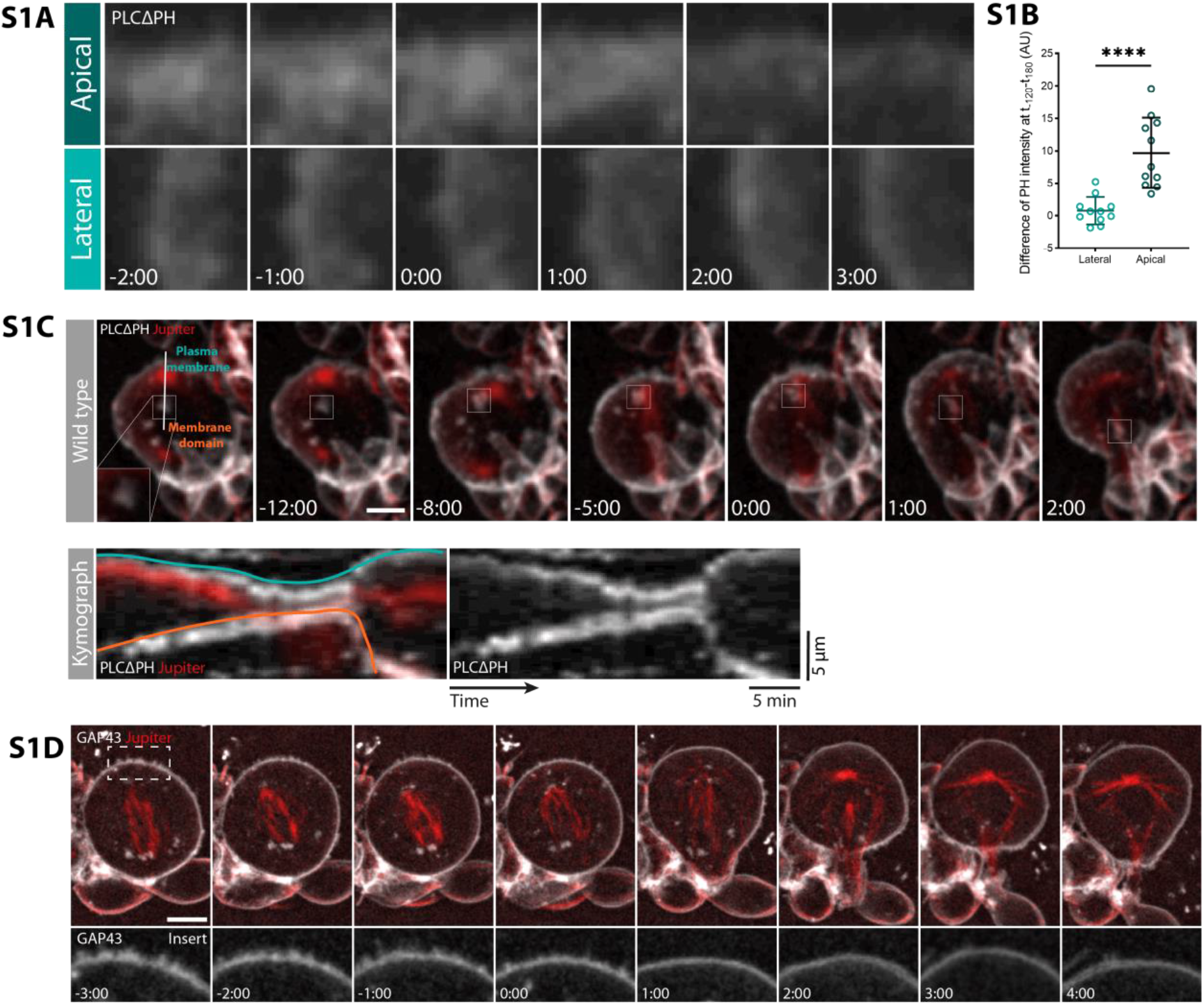
In dividing neuroblasts apical plasma membrane undergoes remodelling dependent on the cell cycle. **A**. Zoom of apical and lateral membrane from wild-type neuroblast expressing UAS-PLCΔPH::GFP. **B**. Plot showing difference of PH intensity between t-120sec and t180sec in both lateral and apical membrane domains. Asterisk (****) denote statistical significance. P ≤ 0.0001 (paired t-test). **C**. Maximum intensity z-projection of dividing neuroblast expressing membrane marker, UAS-PLCΔPH::GFP and a microtubule marker UAS-cherry::Jupiter, both expressed via wor-GAL4/UAS. The white line indicates the position used to generate the kymograph. Insert shows an example of a membrane domain. Kymograph shows the movement of the plasma membrane (blue) and of a more basal PH::GFP-rich membrane domain (orange). **D**. High resolution imaging of neuroblast expressing membrane marker, mCherry::GAP43, driven by sqh promoter. Insert shows membrane protrusions at the apical side of the cell in metaphase (−3:00 to - 1:00). Apical protrusions begin to disappear as cells enter anaphase. Scale bar: 5 μm. Central and error bars: mean and SD.

**Supplemental Figure S2.**
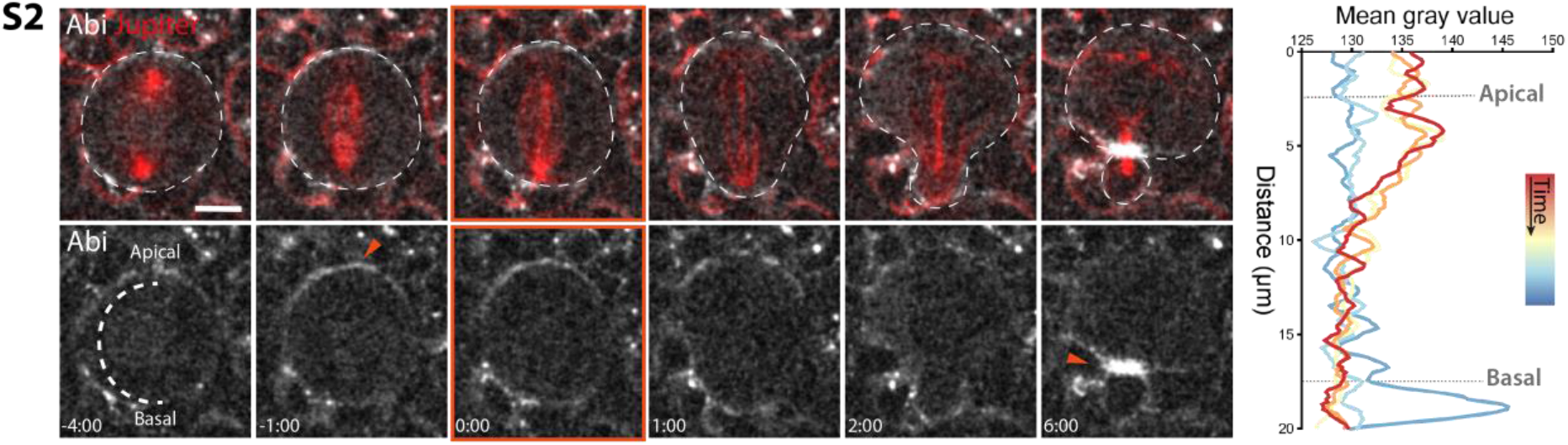
A second component of the SCAR complex, Abi, localizes at the apical side of the neuroblast at metaphase and the furrow at cytokinesis. Super-resolution images of neuroblast expressing ubi-mCherry::Abi. Arrowheads point to Abi localization at the apical cortex in metaphase and at the furrow at cytokinesis. Mean Abi signal intensity was acquired by drawing a line from apical to basal side of the cortex, like depicted at time -4:00, and was plotted in graphs on the right. Scale bar: 5 μm.

**Supplemental Figure S3.**
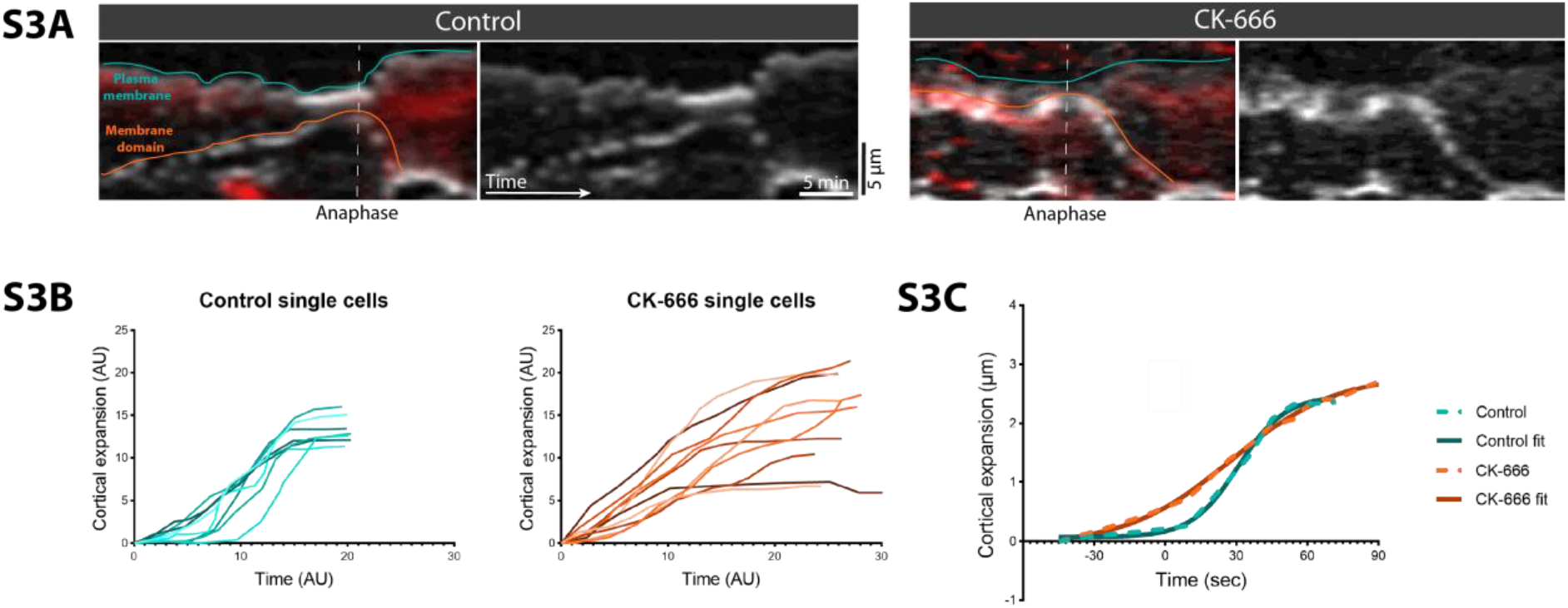
Arp2/3 inhibition does not abolish the movement of membrane domains during neuroblasts cell division. **A**. Kymographs of control (CK-689 treatment) and CK-666 treated cells, expressing PH::GFP and cherry::Jupiter, and showing movement of plasma membrane (blue lines) and membrane domain (orange lines) starting from prophase to cytokinesis. **B**. Single cell tracks of cortical expansion during anaphase in control (CK-689) and CK-666 treated cells. Graph showing mean of cortical expansion and sigmoid curves fitted on top of control and treatment curves.

**Supplemental Figure S4.**
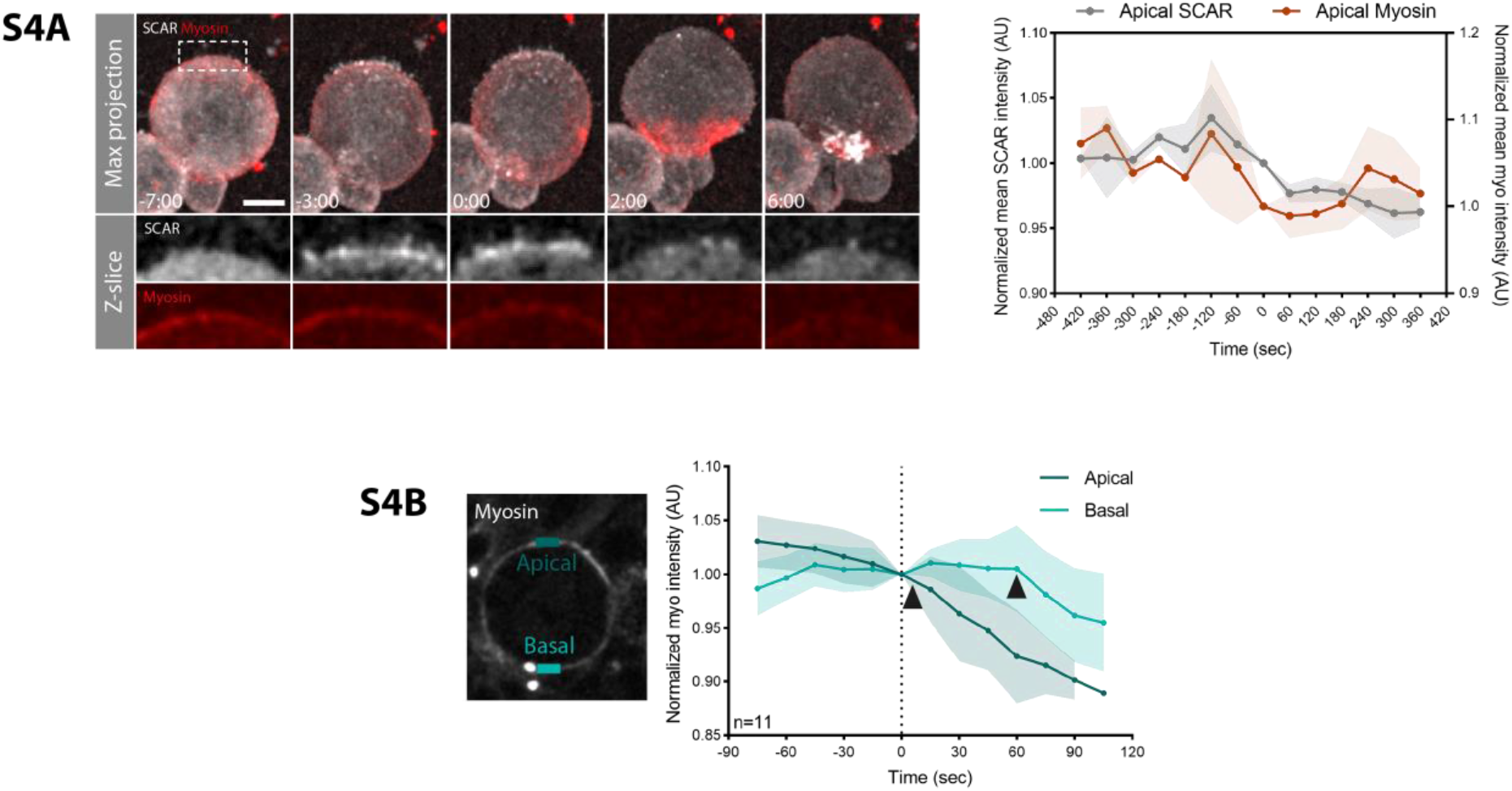
Apical Myosin and SCAR undergo parallel changes in their localisation at the metaphase-anaphase transition. **A**. Super-resolution imaging of dissociated neuroblast expressing SCAR reporter, UAS-SCAR::GFP and a non-muscle Myosin II marker, UAS-Sqh::cherry, which expressions is driven by wor-GAL4. Inserts show apical SCAR and Myosin signals. Graph on the right shows SCAR and Myosin apical intensities during neuroblast division. **B**. Graph shows apical and basal Myosin intensity changes with time, measured from cells expressing Myosin marker Sqh::GFP. Apical and basal rectangles in the cell indicate the areas measured for the graph. Arrowheads mark time points at which Myosin starts to be cleared. Scale bar: 5 μm. Central and error bars: mean and SD.

**Supplemental Figure S5.**
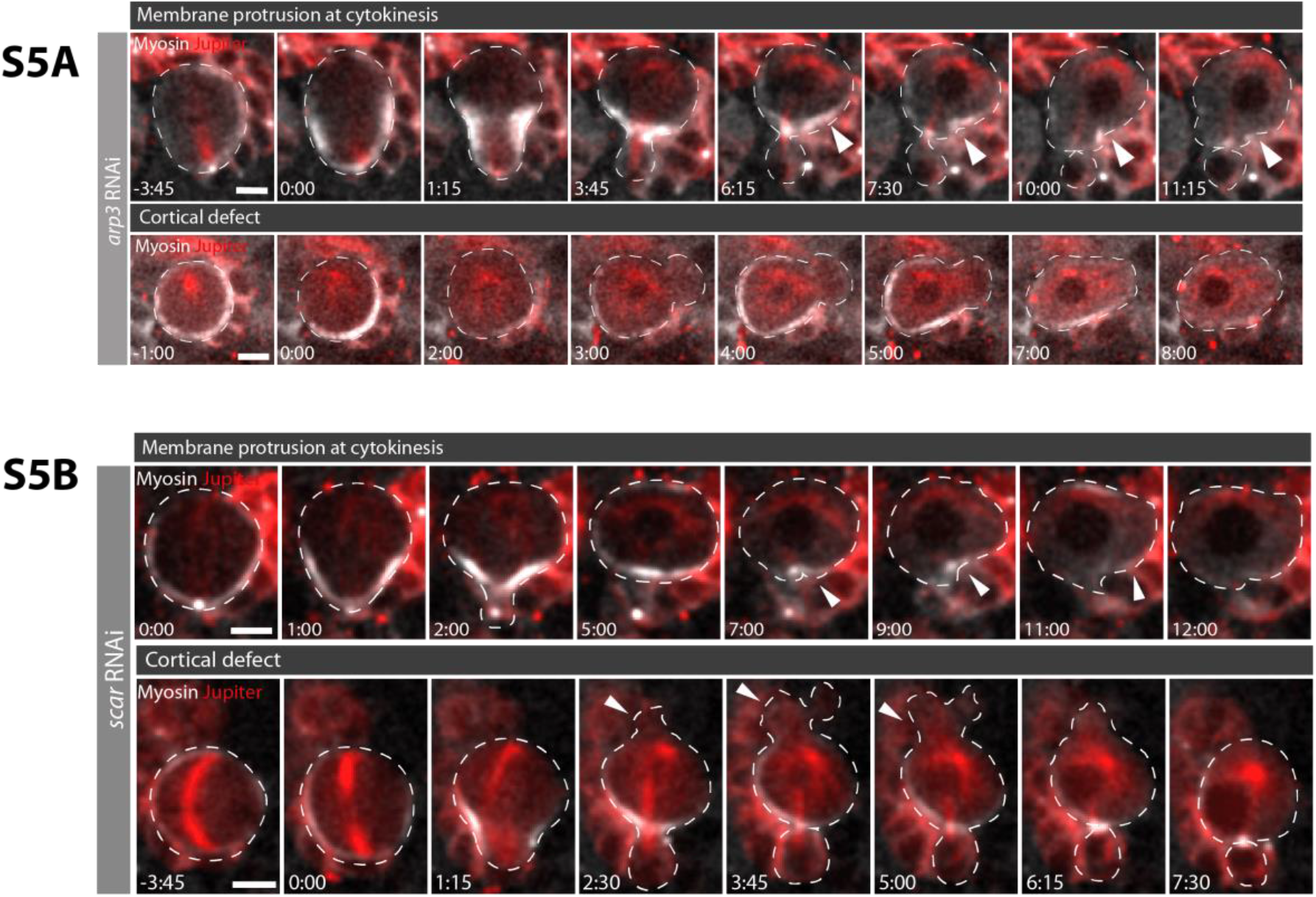
Knock-down of arp3 and scar by RNAi lead to cortical defects and membrane instability at cytokinesis. Time-lapse image of representative neuroblasts expressing Myosin marker Sqh::GFP and microtubule marker cherry::Jupiter. Panels on top show example of membrane protrusion phenotype at cytokinesis. Panels on the bottom show example of cortical defects like blebbing. **A**. Cells expressing dsRNA for arp3 subunit of the Arp2/3 complex. **B**. Cells expressing dsRNA for scar. Scale bar: 5 μm.

## Notes

### Competing Interest Statement

The authors have declared no competing interest.

